# The Proton Resonance Enhancement for CEST imaging and Shift Exchange (PRECISE) family of RF pulse shapes for Chemical Exchange Saturation Transfer MRI

**DOI:** 10.1101/2024.06.19.599565

**Authors:** Zinia Mohanta, Julia Stabinska, Assaf A. Gilad, Peter B. Barker, Michael T. McMahon

## Abstract

**Purpose:** To optimize a 100 msec pulse for producing CEST MRI contrast and evaluate in mice.

**Methods:** A gradient ascent algorithm was employed to generate a family of 100 point, 100 msec pulses for use in CEST pulse trains (‘PRECISE’). Gradient ascent optimizations were performed for exchange rates (k_ca_) = 500 s^−1^, 1,500 s^−1^, 2,500 s^−1^, 3,500 s^−1^ and 4,500 s^−1^ and offsets (Δω) = 9.6, 7.8, 4.2 and 2.0 ppm. 7 PRECISE pulse shapes were tested on an 11.7 T scanner using a phantom containing three representative CEST agents with peak saturation B_1_ = 4 μT. The pulse producing the most contrast in phantoms was then evaluated for CEST MRI pH mapping of the kidneys in healthy mice after iopamidol administration.

**Results:** The most promising pulse in terms of contrast performance across all three phantoms was the 9.6 ppm, 2500 s^−1^ optimized pulse with ∼2.7 x improvement over Gaussian and ∼1.3x’s over Fermi pulses. This pulse also displayed a large improvement in contrast over the Gaussian pulse after administration of iopamidol in live mice.

**Conclusion:** A new 100 msec pulse was developed based on gradient ascent optimizations which produced better contrast compared to standard Gaussian and Fermi pulses in phantoms. This shape also showed a substantial improvement for CEST MRI pH mapping in live mice over the Gaussian shape and appears promising for a wide range of CEST applications.

## Introduction

Chemical exchange saturation transfer MRI has matured as a contrast mechanism that enables the detection of low-concentration contrast agents by applying saturation pulses on resonance with labile protons and measuring the resulting water signal loss based on the chemical exchange transfer of this saturation ^1–4^. Several studies have established the utility of CEST MRI, including measuring mobile protein content of brain tumors for diagnosis ^5, 6^ and discrimination of radiation necrosis from tumor regrowth ^7, 8^, visualizing the perfusion of tumors by glucose and similar sugars ^9–13^, mapping the pH of tumor microenvironment ^14–16^, mapping the pH of the kidneys to detect changes in function ^17, 18^, monitor pH changes after cerebral ischemia, monitor gene expression through CEST reporter genes ^19–21^ among others. The number of adopters of CEST MRI technology has grown further due to new technologies that can improve the speed, selectivity, and sensitivity.^1^

A key issue with CEST MRI is optimization of the pre-saturation of the target exchanging protons in order to maximize CEST contrast. Factors to be considered which influence this contrast include the amplitude, duration and shape (waveform) used for the saturation pulse or pulses. The pre-saturation sequence has to be compatible with scanner hardware, and particularly in human studies, not exceed the maximum allowed specific absorption rate (SAR) or require prolonged periods at maximum power resulting in pulse droop. Optimum pre-saturation parameters will also be dependent on labile proton chemical shifts and exchange rates, as well as relaxation times of both the water resonance and exchangeable protons. Because of scanner hardware constraints, typically CEST MRI experiments are performed with trains of amplitude modulated RF pulses; the shape of the amplitude profile in particular effects the frequency selectivity of the pulse.

Previously, Fermi ^22^, Gaussian ^23^, sinc-Gaussian (‘sg’) convolutions^24^ and Blackman ^25^ pulse shapes have all been used for CEST MRI pulse trains; however, these have not been explicitly optimized previously to maximize CEST contrast. The gradient ascent method of Kuprov, Glaser, and colleagues can be applied to optimize RF pulse design. It has been previously evaluated for designing long RF pulses for paraCEST contrast agent detection in phantoms ^26^. In this paper, the gradient ascent pulse engineering approach is applied to generate a family of ‘Proton Resonance Enhancement for CEST imaging and Shift Exchange’ (PRECISE) generally applicable 100 ms pulses which can be plugged into various CEST pulse sequences, including fast spin echo sequences with a long saturation pulse train preparation module, or steady state CEST sequences with a single pulse for CEST contrast followed by a low flip angle readout module. In particular, ascent optimizations were performed for protons resonating at four labile proton offsets (Δω) and five different exchange rates (k_ca_). Since the most common form of analysis for CEST MRI is to calculate asymmetry between positive and negative offsets from the water resonance (to account for non-specific magnetization transfer effects, and direct water saturation), the cost function used in the simulations was the difference in magnetization between positive and negative offsets. The PRECISE shapes were also compared for how these change for two different relaxation rates and four different mixing times. The 7 most promising PRECISE shapes were tested on three representative diaCEST agents in widespread use in our center, salicylic acid, I45DC-diGlu and iopamidol, in order to evaluate which performed best in terms of contrast generation and pH mapping. Finally, we compared how our best shape performed in comparison to Gaussian pulses for detecting contrast and pH mapping in the kidney after injection of iopamidol into live mice.

## Methods

### Gradient Ascent optimizations

The gradient ascent pulse optimizations were conducted similarly to those described previously by Glaser and colleagues ^26^ using the GRAPE algorithm for optimal control-based design of RF pulses^27^. This divided the pulse into a series of rectangular pieces and modulated the amplitude of each and calculated the evolution of magnetization using a set of 3 pool Bloch equations by numerically solving these using the homogeneous form.

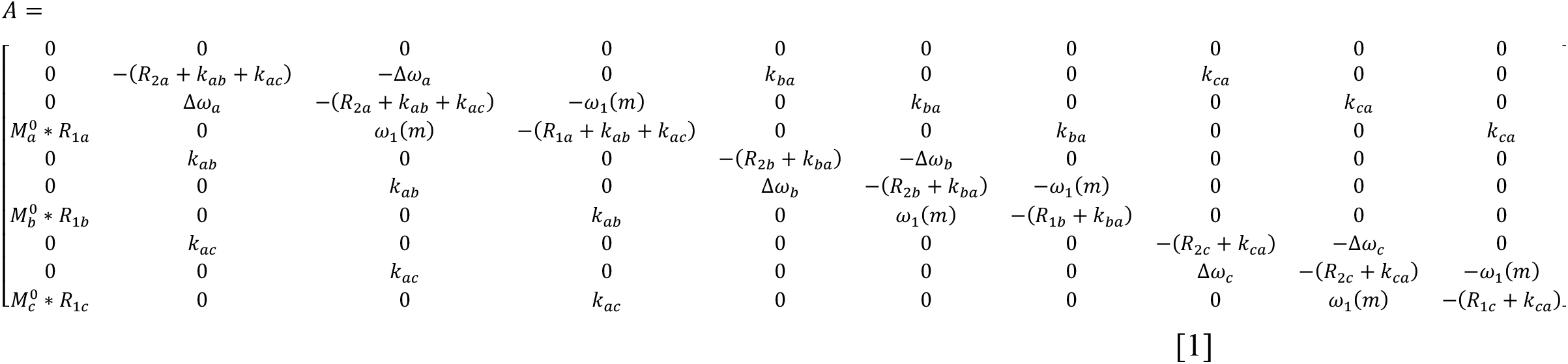

with Δ*ω*_*a*_, Δ*ω*_*b*_, Δω_c_ being the offsets for the three exchanging pools a, b and *c*, ω_1_(m) is the pulse amplitude at time step m, R_1a_ and R_2a_ are the relaxation rates of pool a, M_a_^0^ is the thermal equilibrium magnetization for pool a and k_ab_ is the rate of spin exchange from pool a to pool b.

The pulse segments can then be described by propagators for each segment with for the Mth propagator

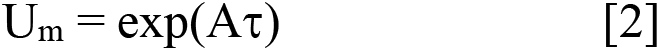

where the length of time for the segment is τ. Given relaxation rates, a control RF and offsets, the propagators for each segment can be calculated independently, and then magnetization evolved through

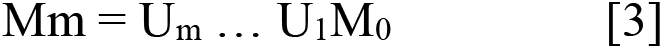

We were interested in optimizing the contrast assuming a water pool (a), a semi-solid pool (b), and one CEST contrast agent pool (c) and employed the Bloch equations with chemical exchange terms present. We assume there will not be any side exchange between semi-solid and CEST pools b and c. The initial control was set to a train of Gaussian shaped pulses, and the figure of merit or cost function to assess the performance of the pulses is:

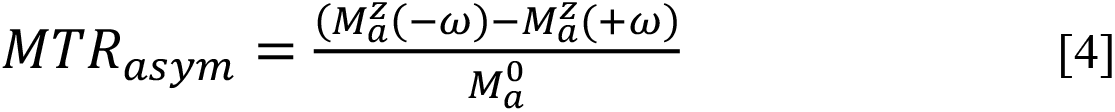

where 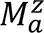 is the z magnetization of water at equilibrium. The gradient is calculated based on changes with respect to the control.

#### Optimization Parameters

The relaxation and semi-solid pool parameters were set based on previous studies on CEST and magnetization transfer contrast. ^28–31^ The parameters include: semi-solid pool concentration (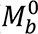) = 1,554 mM, CEST concentration (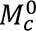) = 50 mM, B_0_ field = 11.7 T (500 MHz), k_ba_ = 20 s^−1^, k_ca_ = 500 – 6,500 s^−1^, T_1a_ = 2 sec, T_1b_ = 1 sec, T_1c_ = 1 sec, T_2a_ = 60 msec, T_2b_ = 10 μsec, T_2c_ =100 msec. We employed a 1 msec timestep for optimizing the 100 msec pulses (100 steps per 100 msec pulse) and a delay between of 1 to 100 msec between the pulses with a total of 2000 total steps for this pre-saturation pulse train including the pulses and delays. The pulses were also set to be identical for each of the up to 19 pulses in the pulse train.

#### Simulation of magnetization plots

To characterize how the individual pulse shapes would perform as inversion pulses and compare the performance to the Gaussian, sg100 and Fermi pulses, we used the ‘Shape Tool’ embedded within Bruker Topspin software version 3.5 to simulate the frequency dependence of the inversion. We assumed a pulse duration = 100 msec, with approximately 4 μT peak power, however the exact nutation was set to an odd integer of 180°. All pulses were 100 points, and the exact nutation angle reported in **Table 2**. R_BW_ as described in de Graaf et al ^32^, was calculated as Bandwidth (kHz) × pulse length (msec) measured at M_z_/M_0_ = 0.0. The pulse integral was calculated by integrating and dividing by a rectangular pulse with 100% amplitude across all 100 points. The pulse root mean square (RMS) integral was calculated by integrating the square of the pulse over the 100 points and then taking square root of the result. The Side Band Ripple (SBR) mean and variance were calculated from frequencies of −75 to −150 Hz from the center.

#### In vitro Phantom Experiments for testing CEST performance of pulses

To evaluate the pulse shapes, a phantom was prepared consisting of 5mm NMR tubes, with one filled with 0.01 M phosphate-buffered saline (PBS) as the negative control, and the others containing three CEST agents: 1) salicylic acid, 2) I45DC-diGlu and 3) iopamidol. Tubes were prepared with agents in phosphate buffered saline (PBS) at concentrations of 50 mM and the pH adjusted to 7.0. In addition, tubes were prepared containing 1% agarose by heating agarose at 50°C for 10 minutes and while cooling adding 50 mM of each agent and titrating the resulting solution to pH 7.0.

#### In vitro phantom experiments for CEST protocol optimization for pH mapping

To evaluate the CEST performance using one of the optimized pulses and one of the standard pulses, a phantom consisting of 10 5 mm NMR tubes, with one filled with 0.01 M phosphate-buffered saline (PBS) as the negative control and the other tubes consisted of 40 mM iopamidol solutions titrated to pH 6.0, 6.2, 6.4, 6.6, 6.8, 7.0, 7.2, 7.4 and 7.6. For CEST data, we employed the RARE sequence with specific parameters: TE/TR: 20/6000 msec, matrix size: 64 × 64, slice thickness: 1.5 mm, and RARE factor: 32. Saturation parameters were peak B_1_ = 4 μΤ, t_pulse_ = 100 msec, t_mix_ = 1 msec and total t_sat_ = 2.9 sec.

### In vivo experiments

Experiments were performed in accordance with Institutional Animal Care and Use Committee guidelines. MR images were acquired on a Bruker Biospec 11.7 T scanner, using a 72 mm body coil for transmission and an 8-channel phase-array coil for reception. Two groups, each consisting of 3 C57Bl/6 mice, were tested for the PRECISE 9.6 2500 pulse and the standard gaussian pulse. The tail vein was cannulated for the administration of iopamidol. The mice (12-16 weeks, male; National Cancer Institute, Frederick, MD) were anesthetized with 2 % isoflurane and kept warm with a heating pad. The animals were securely positioned on a transceiver body coil and subjected to anesthesia with 0.5%–2% isoflurane. Continuous monitoring ensured stable respiratory rates, while the mice’s body temperature was maintained at 37°C using a water-heated animal bed. Imaging comprised 25 axial and 7 coronal contiguous slices obtained through a multi-slice T2w RARE sequence with specific parameters: TE/TR: 20/4000 msec, matrix size: 128 × 128, slice thickness: 1.5 mm, and RARE factor: 8. A single 1.5 mm thick axial slice, centered on both kidneys, was selected for CEST imaging. B0 inhomogeneity maps were generated utilizing the water saturation shift referencing (WASSR) approach, involving acquisition at 42 frequency offsets between −1.5 and 1.5 ppm, employing 10 saturation pulses with a duration of 300 msec and an inter-pulse delay of 10 μsec, with B1 = 1.2 μT. Dynamic CEST experiments utilized a saturation module comprising 29 RF pulses, each lasting 100 msec with an inter-pulse delay of 1 ms and an amplitude of 4 μT. The readout parameters consisted of TE/TR = 3.49/5000 msec, matrix size: 32 × 32, FOV: 28 × 20 mm^2^, slice thickness: 1.5 mm, and RARE factor: 33. A total of 234 CEST images were acquired, comprising 10 images at 40 ppm and 112 sets of images at 4.3 and 5.5 ppm. 0.15 mL of iopamidol (0.370 M in saline) was administered over 1 min, 3 minutes after the commencement of CEST data acquisition. The entire CEST scan duration amounted to 19 min 30 sec.

### In vitro pH calibration

Post-processing and pH calculations for our phantom were conducted using MATLAB (MathWorks, USA). Following previously established methods, ^33^ these computations involved determining the saturation transfer (ST) ratio at 4.2 and 5.5 ppm. Pixel-by-pixel Z-spectra were computed from the CEST data, with B_0_ inhomogeneity corrections applied using the WASSR dataset and spline interpolation technique, as outlined in previous studies. The ST value, calculated as (1 − M_z_/M_0_) pixel by pixel, was utilized to generate ST maps corresponding to the CEST peaks at 4.2 and 5.5 ppm, respectively. Subsequently, regions of interest (ROIs) were delineated, and the mean ST ratio versus pH plot was generated and fitted to a polynomial to establish a pH calibration curve. The pH for each pixel in the phantom was determined using this calibration equation.

### In vivo Post-Processing and CEST data analysis

Custom-written MATLAB (Mathworks, Natick, MA, USA) scripts were utilized for image post-processing and dynamic CEST data analysis. ^33^ Regions of interest (ROIs) were drawn across entire cross section of both kidneys to obtain the signal intensity maps and pH maps. A moving average filter (implemented using the MATLAB function filter) with a span of 2 was applied at each frequency offset to mitigate motion-induced signal fluctuations. Region of interest (ROI)-based time-course curves of signal enhancement were derived by plotting the percentage change in CEST-prepared signal intensity at 4.2 ppm post-iopamidol injection (M_z,post_) relative to the averaged signal from seven pre-injection images (M_z,pre_):

[eqn1]

For pH mapping, post-injection magnetization transfer ratio (MTR) at 4.2 and 5.5 ppm was quantified from three averaged images acquired at peak enhancement time. Pre-injection MTR maps were averaged and subtracted from post-injection MTR images to eliminate endogenous CEST signals. Subsequently, renal pH values were determined by calculating the concentration-independent saturation transfer ratio:

[eqn1]

and using pH calibration curves obtained in the *in vitro* phantom study.

### Statistical significance

In this study, we employed the Mann-Whitney test to assess the statistical significance of differences between the two groups of mice (n=5) employing the optimized pulse and standard pulse. The Mann-Whitney test, also known as the Wilcoxon rank-sum test, is a non-parametric test suitable for comparing independent groups with non-normally distributed data and provides robust results even with small sample sizes. P<0.05 was considered to be statistically significant.

## Results

Typical agents that are used in our center are shown in **Fig 1A** and possess labile protons resonating from 1 to 10 ppm from water. These agents also possess a range of chemical exchange rates (k_ca_) from 400 s^−1^ to 14,000 s^−1^, as described in previous publications, which allows a thorough assessment of how shift and exchange properties impact the ideal pulse shapes for producing CEST contrast. It was first important to determine if and when convergence would occur for the range of k_ca_ and Δω using the pulse scheme shown in **Fig. 1B**, imposing the condition that all pulses in the train have the same shape and amplitudes. Most studies published so far employ the Gaussian, sg100 or Fermi shapes shown in **Fig. 1C**. It is interesting to see how close these shapes are to optimized pulses when considering the other sources of water signal loss during saturation pulse trains, notably direct saturation of water and conventional magnetization transfer.

**Figure 1.**
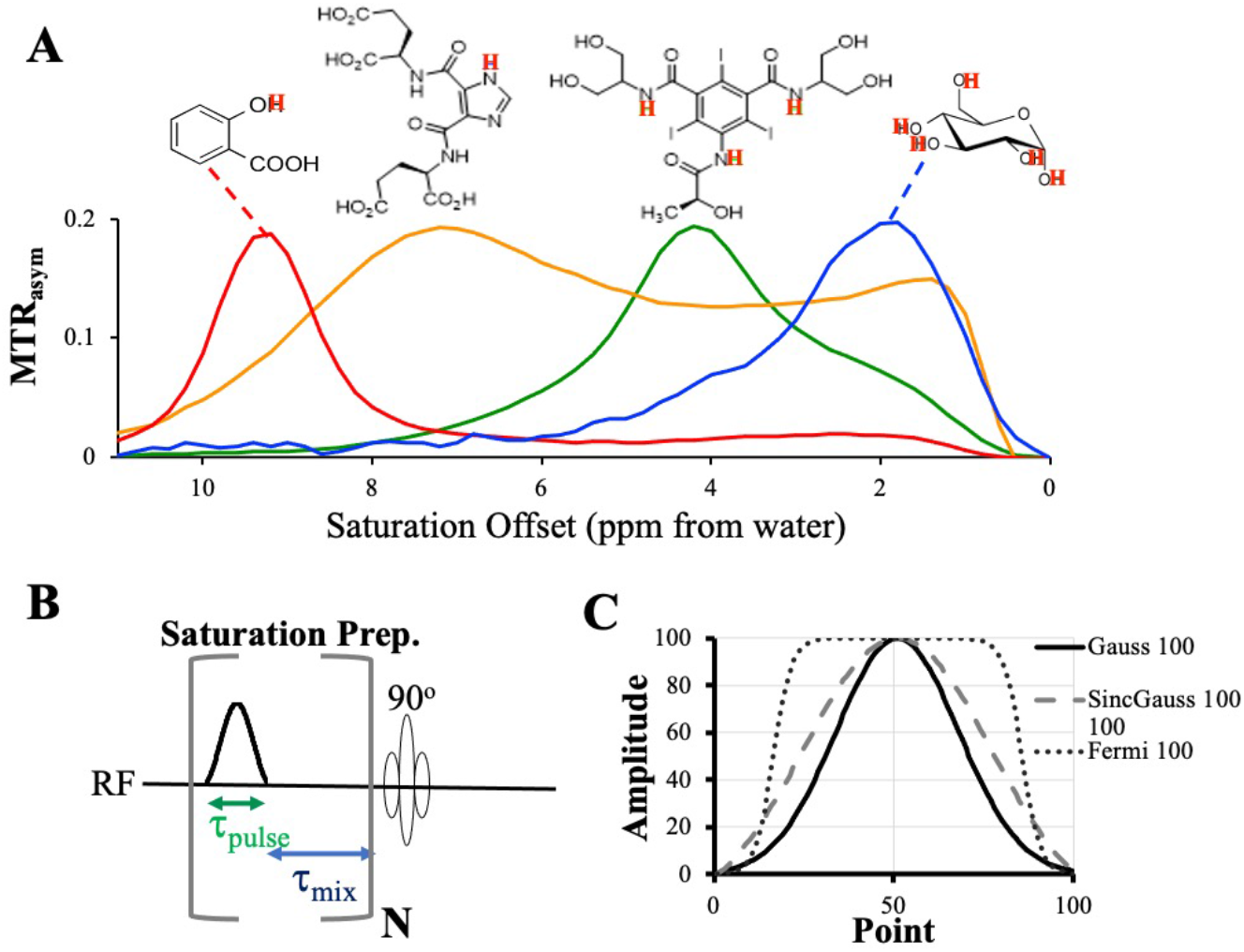
Optimization scheme for representative CEST agents. **A)** CEST MTR_asym_ spectra and chemical structures of the 4 representative agents included in this study: salicylic acid, I45DC-diGlu, iopamidol and d-glucose. Conditions: CW saturation with B_1_ = 2.6 μT, T_sat_ = 2 sec, TR = 5 sec; **B)** Pulse scheme employed for optimizations with pulse length and mixing time fixed; C) Common pulse shapes employed for CEST pulse trains including Gaussian, convolution of sinc and Gaussian (1.5% cutoff) and Fermi pulses.

We first had to determine how well our optimizations would converge using the cost function in Eqn. [4] with our initial guess when τ_pulse_ = 100 msec, τ_mix_ = 10 msec for the saturation prep shown in **Fig. 1B** and relaxation, and MT contrast parameters set to mimic kidney tissue. We started our optimizations with Δω = 9.6 ppm, 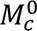 = 50 mM as this is our furthest shifted agent so the direct saturation of water will be the smallest for this shift and ∼ 50 mM is an achievable concentration in the kidneys for multiple agents. **Fig. 2** displays our results, which indicate that for k_ca_ = 500 s^−1^, 1,500 s^−1^ and 2,500 s^−1^ additional iterations do not result in a tangible change in pulse shape by ∼6,500 steps (**Fig. 2A-C**), while for k_ca_ = 3,500 s^−1^ this requires 9,500 steps (**Fig. 2D**) and for k_ca_ = 4,500 s^−1^ this requires 19,500 steps (**Fig. 2E**). Unfortunately for k_ca_ = 6,500 s^−1^ there are still substantial changes in shape even after 39,000 steps (**Fig. 2F**), presumably based on the very small changes in contrast which might occur for a wide range of pulse shapes. We also verified how the simulations would converge for the lowest shifted labile proton Δω = 2.0 ppm, with again ∼9,500 steps to achieve convergence for k_ca_ = 3,500 s^−1^ (**Fig. 2G**) and ∼19,500 steps for k_ca_ = 4,500 s^−1^ (**Fig. 2H**). From these simulations it is also clear there are difficulties optimizing pulses for k_ca_ ≥ 4,500 s^−1^. We also found similar convergence results for t_mix_ = 1 msec, which is expected as the contrast will be larger as the delay is decreased. Based on this, we went ahead and optimized pulses for labile proton offsets for all three agents we planned on evaluating as part of this study with our results shown in **Fig. 3**. What can be seen quite clearly is that for the faster k_ca_ = 3,500 and 4,500 s^−1^, as the Δω increases the pulses grow shorter. Indeed by Δω = 9.6 ppm, k_ca_ = 4,500 s^−1^ the pulse amplitude doesn’t increase above zero until about point 20 for t_mix_ = 1 msec, while for Δω = 2.0 ppm, k_ca_ = 4,500 s^−1^, the pulse immediately rises to 40% of its peak height in the first few points. There is a general increase in oscillations as the offset increases that can be observed as well. However, this figure also demonstrates that the optimal shapes are fairly similar for 3 of the 4 offsets when k_ca_ is held the same, with the shapes for Δω = 2.0 ppm being different presumably due to the much larger direct saturation of water when irradiating at this frequency. In summary, it is possible to obtain optimized pulse shapes by using these preparation parameters and the gradient ascent algorithm. There are similar shapes that the simulations settle upon across the range of labile proton offsets evaluated. In addition, this first search of optimal 100 msec CEST pulse shapes produced candidates which were quite different from the standard shapes in **Fig. 1C**, with the optimized shapes changing as a function of k_ca_ and Δω.

**Figure 2.**
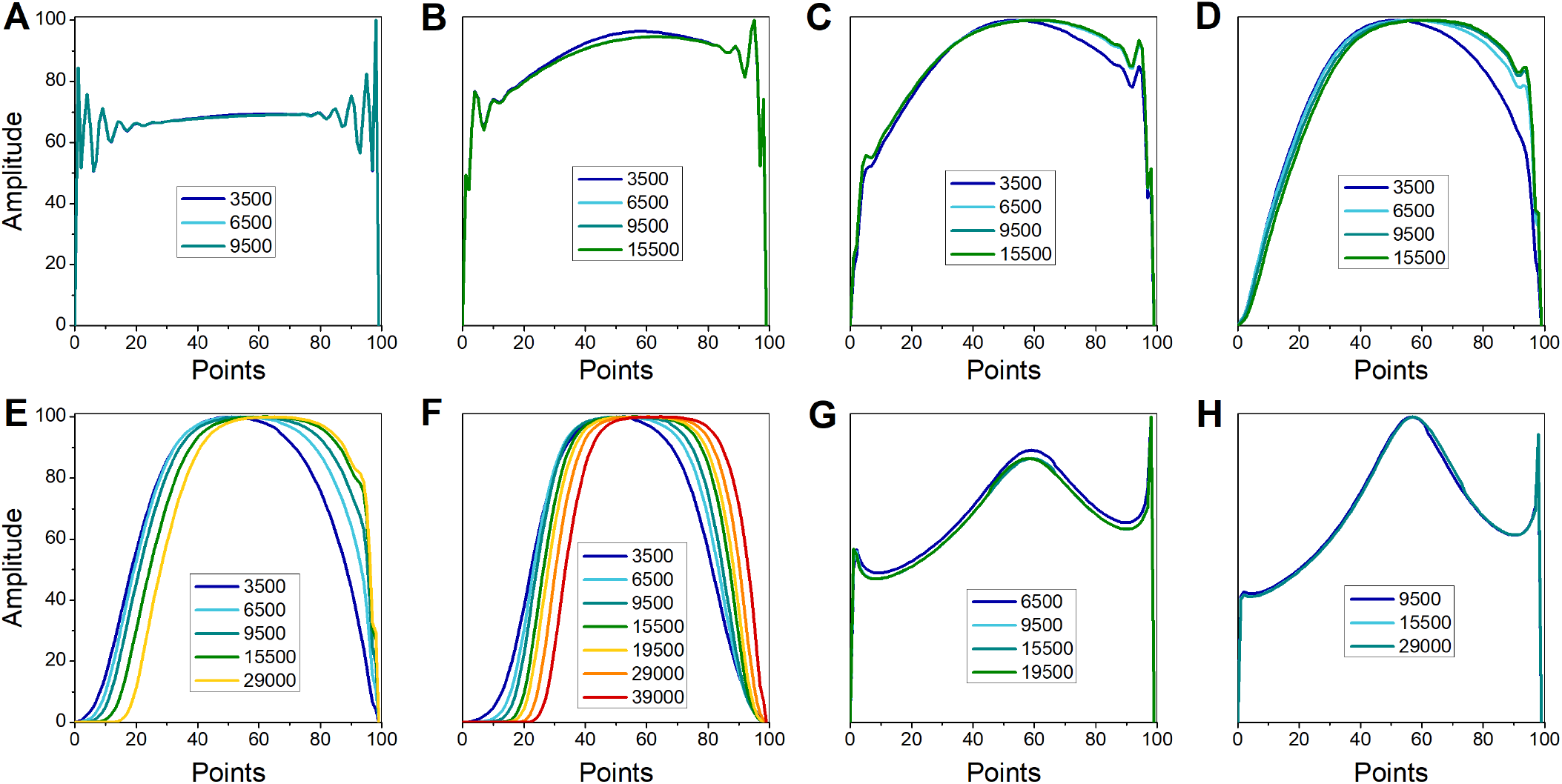
Pulse shapes for a given number of steps using the gradient ascent method for determining convergences with **A-F** Δω = 9.6 ppm and **G-H** Δω = 2.0 ppm and number of iterations for the gradient ascent algorithm listed in the individual plot legends. These simulations employed 1_pulse_ = 100 msec, 1_mix_ = 10 msec and 18 pulses. For Δω = 9.6 ppm, we tested convergence for k_ca_ = **A)** 500 s^−1^; **B)** 1,500 s^−1^; **C)** 2,500 s^−1^; **D)** 3,500 s^−1^; **E)** 4,500 s^−1^; **F)** 6,500 s^−1^. For Δω = 2.0 ppm, we tested convergence for k_ca_ = **G)** 3,500 s^−1^; **H)** 4,500 s^−1^;

**Figure 3.**
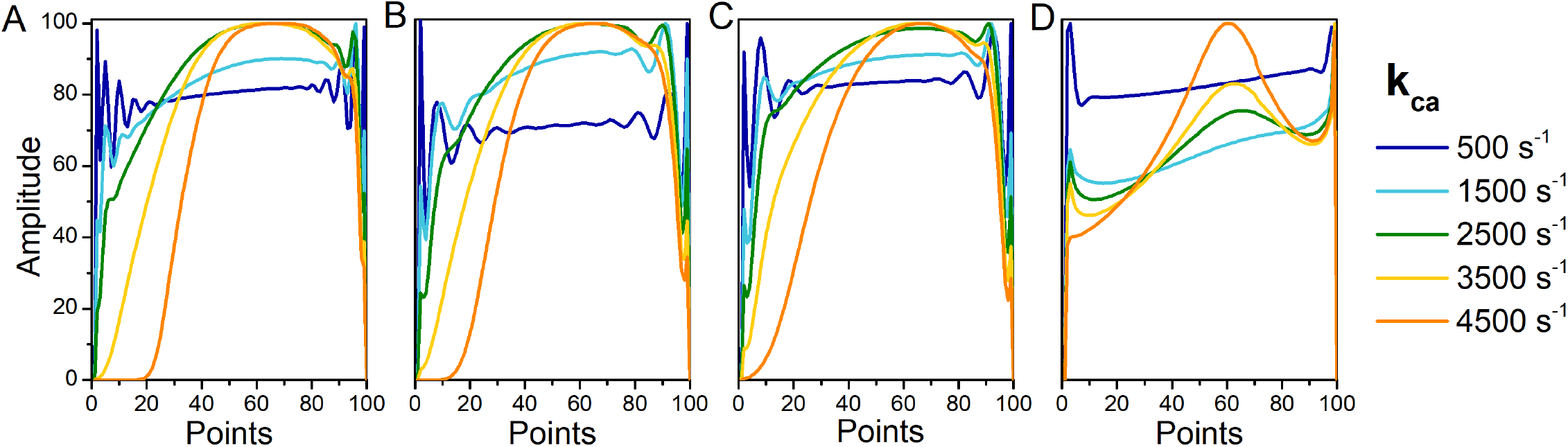
Optimized pulse shapes as a function of ι1ω and k_ca_. A) 9.6 ppm; B) 7.8 ppm; C) 4.2 ppm; D) 2.0 ppm. These simulations employed 1_mix_ = 1 msec and 19 pulses.

To conduct a more exhaustive search for candidate pulse shapes, we performed simulations with variations in 1_mix_ = 1 to 100 msec with the results are shown in supplementary figure **Fig. S1**. This variation in 1_mix_ was chosen to investigate the shapes for pulse trains with as low as a 50% duty cycle, which is a particularly common choice for clinical 3 T scanners. We didn’t separately calculate for both 7.8 and 9.6 ppm based on the similarity in the shapes seen in **Fig. 3**. As can be seen as 1_mix_ increases, the pulse shapes increase in width for Δω = 9.6 ppm and 4.2 ppm, with the central region growing for Δω = 2 ppm. The average amplitude also increases as 1_mix_ increases. Furthermore, we considered how water longitudinal relaxation time would impact pulse shape with the results shown in supplementary figure **Fig. S2.** Changes in relaxation time did not have such a big impact on pulse shape for the three offsets considered. In summary, we generated a family of shapes that appear promising for CEST imaging.

Next, the performance of these optimized pulses were evaluated on the phantom which included 3 of our standard CEST contrast agents with a range in Δω and exchange rates and number of labile protons. We list the Δω and k_ca_‘s for pH values typically seen physiologically in **Table 1**. The 7 most promising PRECISE shapes were selected for testing based on the approximate values expected for the agents and are shown in **Fig. 4**.

**Figure 4.**
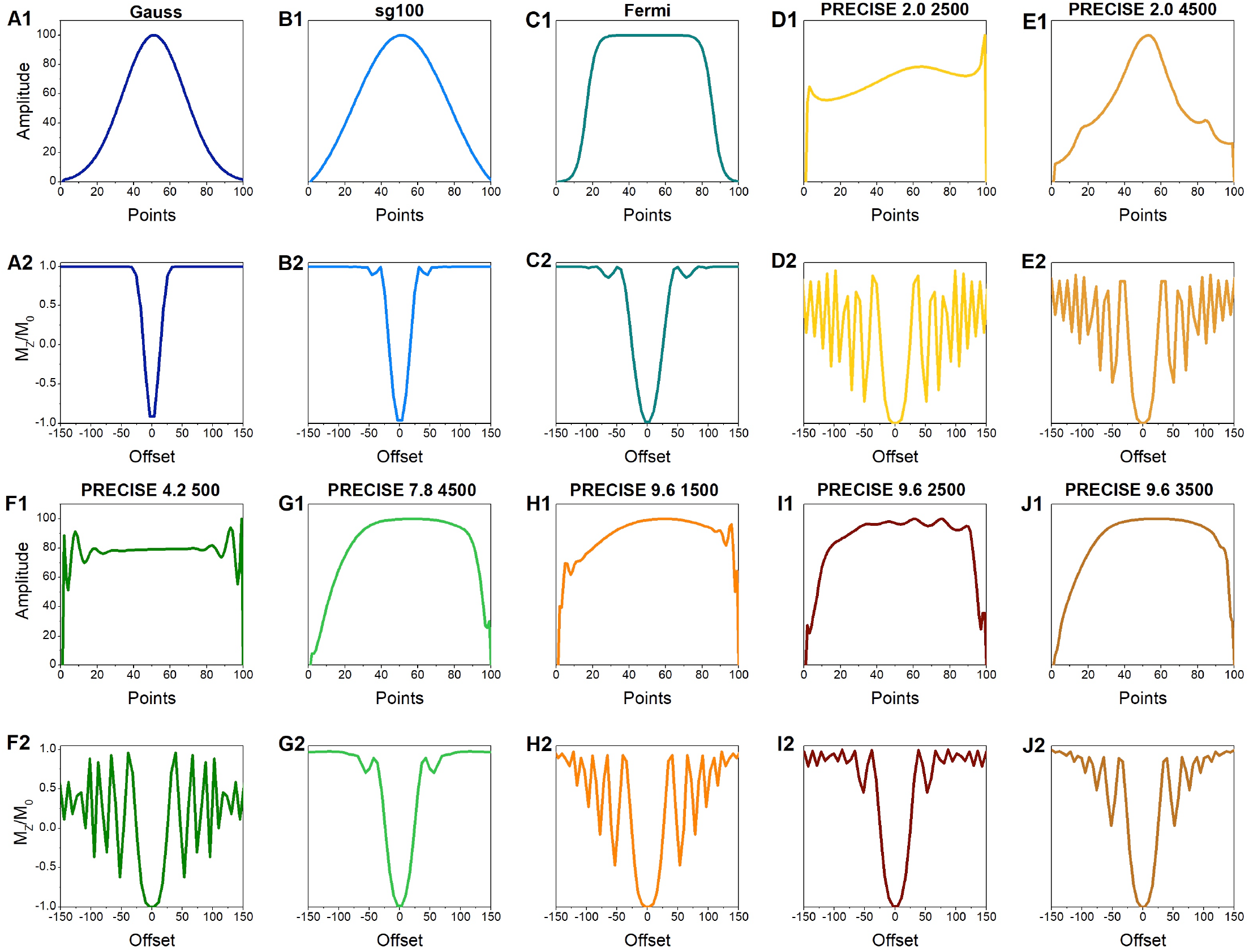
Selected optimized pulse shapes and their inversion profiles.

**Table 1.**
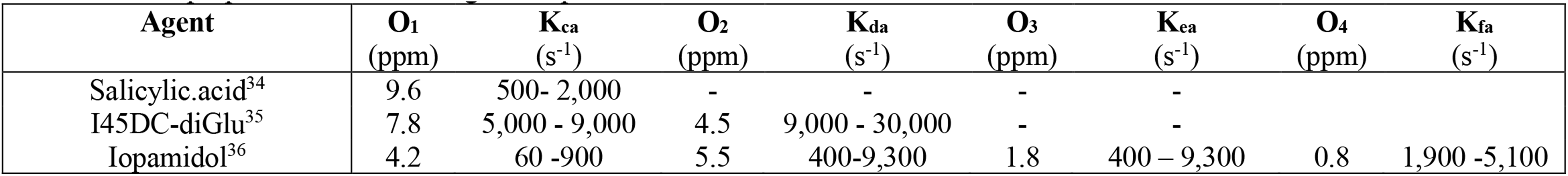
CEST properties for selected agents at pH 6.5 - 7.2.

To understand the frequency selectivity of these pulses for an appropriate saturation field strength for CEST contrast, we calculated the rms B_1_, number of nutations, inversion bandwidth, pulse integral, pulse RMS integral and stop band ripples and display these in **Fig. 4D-J** and **Table 2**. As expected, the gaussian shape inverts magnetization has the smallest nutation angle, pulse length-bandwidth product (R_BW_) and also a low pulse integral, with the sg100 pulse displaying increase, and the Fermi pulse displaying a further increase in these parameters. All of the PRECISE shapes have larger nutation angles, R_BW_ and pulse integrals than the Gaussian, sg100 and Fermi pulses, although the PRECISE shapes also display higher side band ripples which may degrade asymmetry contrast. We named these pulses by which Δω and k_ca_ were used for optimization, so PRECISE 9.6 2500 was optimized using Δω = 9.6 ppm, k_ca_ = 1,500 s^−1^. Of the PRECISE pulses, the PRECISE 9.6 2500 and PRECISE 7.8 4500 pulses display the smallest side band ripples. These two PRECISE shapes appear particularly promising as CEST pulses.

**Table 2.** Characteristics of the individual pulses.

| Pulse | Nutation angle<sup>1</sup> | $R_{BW}$<sup>2</sup> | Pulse integral | RMS integral | Side Band Ripples (SBR) | |
| --- | --- | --- | --- | --- | --- | --- |
|  |  |  |  |  | Mean | Variance |
| Gauss | 2700 | 2.888 | 0.41 | 0.55 | 0.99 | 6.41E-07 |
| sg100 | 3420 | 3.6 | 0.54 | 0.64 | 0.98 | 8.84E-04 |
| Fermi | 4140 | 5.225 | 0.69 | 0.81 | 0.97 | 2.1E-03 |
| PRECISE 9.6 1500 | 4860 | 5.59 | 0.87 | 0.89 | 0.61 | 0.18 |
| PRECISE 9.6 2500 | 5220 | 5.234 | 0.84 | 0.87 | 0.85 | 0.02 |
| PRECISE 9.6 3500 | 4500 | 5.234 | 0.82 | 0.87 | 0.78 | 0.07 |
| PRECISE 7.8 4500 | 3780 | 5.07 | 0.64 | 0.85 | 0.93 | 0.01 |
| PRECISE 4.2 500 | 4860 | 5.305 | 0.77 | 0.78 | 0.27 | 0.19 |
| PRECISE 2.0 2500 | 4140 | 4.816 | 0.68 | 0.7 | 0.29 | 0.29 |
| PRECISE 2.0 4500 | 4140 | 4.754 | 0.69 | 0.58 | 0.9 | 0.19 |

<sup>1</sup> Nutation angle is rounded up to closest odd integral multiple of 180° to obtain complete inversion
<sup>2</sup> $R_{BW}$ = Bandwidth (kHz) × pulse length (ms) measured at $M_z/M_0=0.0$
<sup>3</sup> RMS (Root mean square) integral was calculated by integrating the square of the pulse over time period and then taking square root of the result.

To evaluate the 7 PRECISE shapes for their ability to produce CEST contrast within a train of pulses, we performed a CEST experiment using a 3 sec long train of each of these shapes with 1_pulse_ = 100 msec, 1_mix_ = 1 msec, max B_1_ = 4 μT and measured the CEST contrast for each tube. We first applied the pulses optimized for 9.6 ppm and 7.8 ppm to SA and I45DC-diGlu, our largest labile proton shift agents, with the resulting MTR_asym_ spectra and peak contrast values for each pulse shown in **Fig. 5**. For both agents, we observed maximal CEST contrast using the PRECISE 9.6 2500 pulse, with a particularly large improvement over Gaussian pulses but also a pronounced improvement over the SG100 pulse. A larger increase in contrast is seen for I45DC-diGlu dissolved in PBS compared to 1% agarose, which will also have longer T_1_ relaxation times (T_1_ I45DC in PBS = 3.07 sec, T_1_ I45DC in 1% agarose = 2.08 sec). As can be seen, the PRECISE 9.6 2500 pulse shows outstanding performance for both of these agents with a factor of ∼ 2.7 improvement over Gaussian pulses and 1.3 improvement over Fermi pulses. We further evaluated our pulse shapes on a 2^nd^ phantom containing iopamidol in both agarose and PBS with the results shown in **Fig. 6**. Again, the PRECISE 9.6 2500 produced the strongest contrast at 4.2 ppm, even over the PRECISE 4.2 500 pulse. Based on these phantom experiments, we decided to evaluate the PRECISE 9.6 2500 pulse for *in vivo* pH mapping.

**Figure 5.**
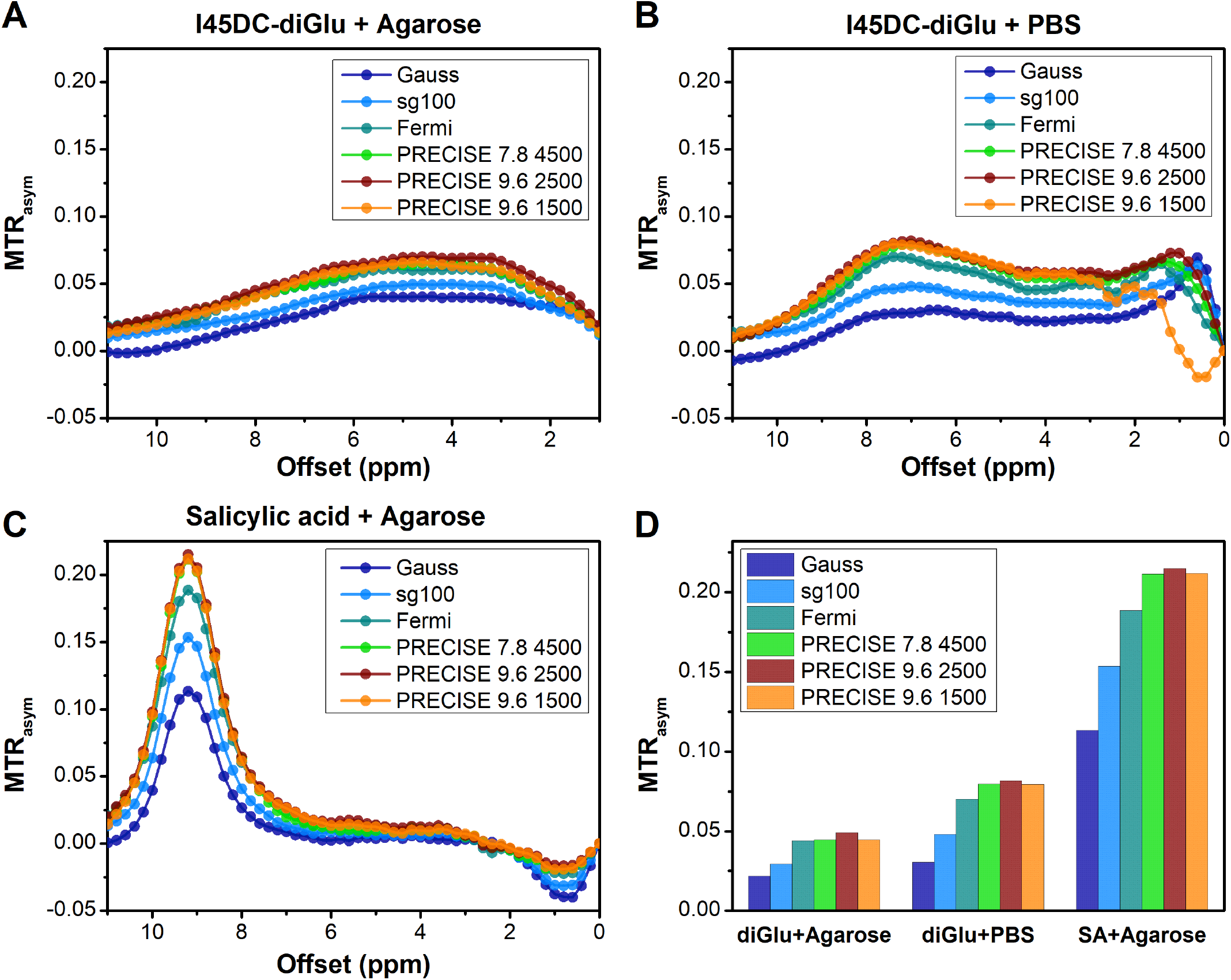
Experimental results for PRECISE pulses on SA and I45DC-diGlu phantoms. **A**) is the MTR_asym_ spectra for I45DC-diGlu in 1% agarose;. **B**) MTR_asym_ spectra for I45DC-diGlu in PBS; **C**) MTR_asym_ spectra for SA in 1% agarose; **D**) Bar plot comparing the performance at the peak contrast frequencies.

**Figure 6.**
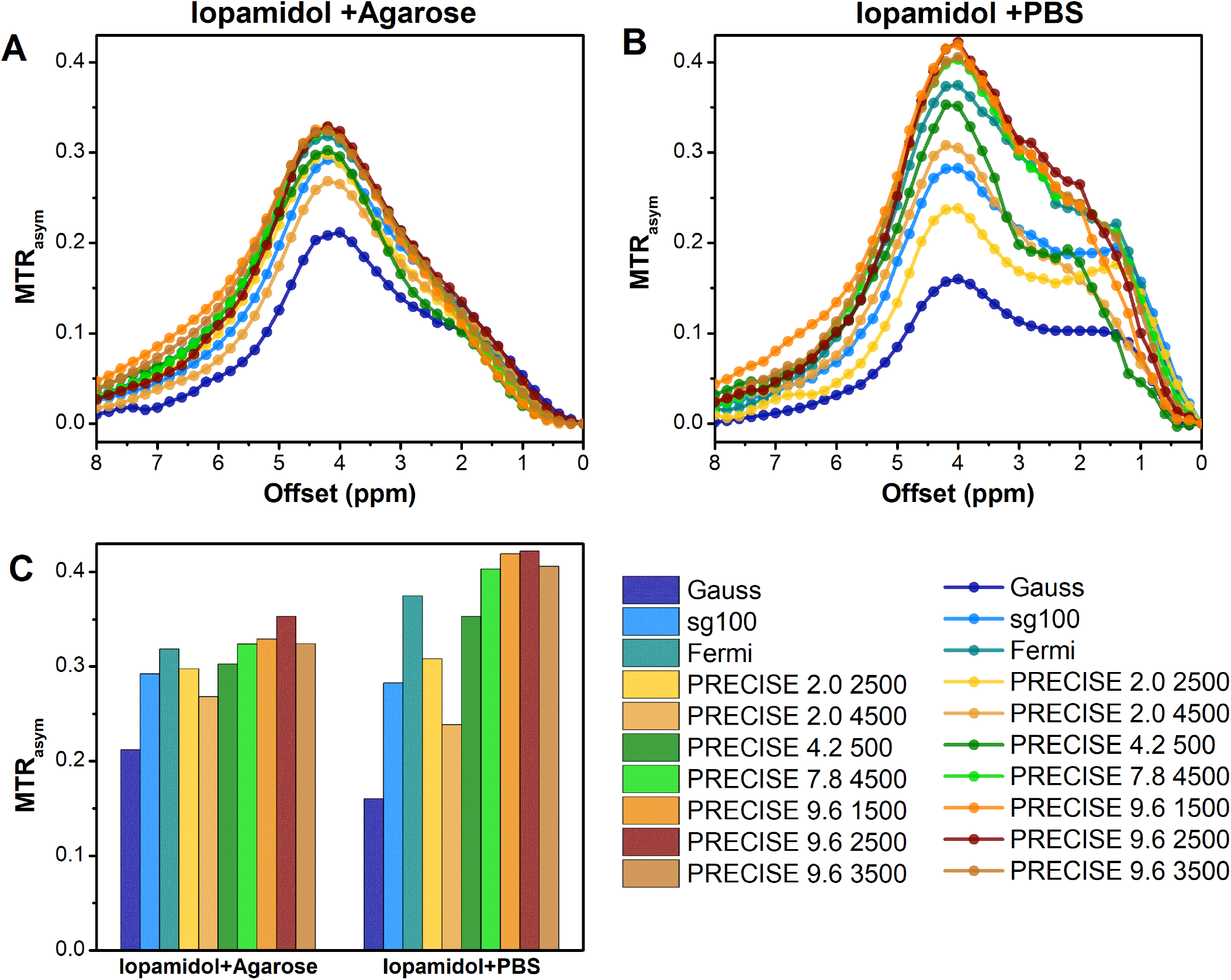
Experimental results for pulses optimized for 4.2 ppm on an iopamidol phantom. **A**) is the MTR_asym_ spectra for Iopamidol in 1% agarose;. **B**) MTR_asym_ spectra for Iopamidol in PBS; **C**) Bar plot comparing the performance at the peak contrast frequencies.

We initially evaluated how the PRECISE 9.6 2500 pulse performed for pH mapping on a phantom consisting of iopamidol in PBS titrated over a series of pH values and compared the contrast to that produced by a standard gaussian shape (**Fig. 7**). A saturation B_1_ = 4 μT was used based on wanting to balance contrast with peak width which is similar to that used on other pH studies^17^. As is shown, the CEST contrast at 4.2 ppm and 5.5 ppm both increase with increasing pH up until pH 7.0, where the CEST contrast value at 5.5 ppm decreases. The highest values of CEST contrast at 4.2 ppm were 38.1 % and 22 % for PRECISE 9.6 2500 and Gaussian respectively at pH 7.4. The highest values of CEST contrast at 5.5 ppm were 21.2 % and 11.1 % for PRECISE 9.6 2500 and Gaussian respectively at pH 7.4. The variation of experimental ST ratio at different pH values was given by pH = p1 × (ST ratio)^3^ + p2 × (ST ratio)^2^ + p3 × (ST ratio)1 + p4. For the Gaussian pulse, the coefficients with 95% confidence bound for the third polynomial fit were p1=-3.718, p2 = 7.454, p3 = −5.962 and p4 =8.652. The ST values fitted for pH values range of 6.2 to 7.4 and varied from 1.06 at pH 6.2 to 0.31 at pH 7.4. For the PRECISE 9.6 2500 pulse, the coefficients with 95% confidence bounds for the third polynomial fit were p1=-0.8318, p2 = 2.394, p3 = −3.073 and p4 =8.288. The coefficient for determination, R^2^ and root mean square error, RMSE for both the fittings were 0.99 and 0.02, respectively. The ST values fitted for pH values range of 6.2 to 7.4 and varied from 1.53 at pH 6.2 to 0.39 at pH 7.4. As is seen from **Fig. 7**, the range of ST values for the PRECISE 9.6 2500 increased by 50% over the range of ST values for the Gaussian pulse. The slope of the pH vs ST curve for PRECISE 9.6 2500 was also lower than the Gaussian slope (−0.77 compared to −1.03). This gradual slope produced by the PRECISE 9.6 2500 pulse and broader range for ST values allows for better pH determinations compared to the Gaussian pulse, as can be seen by the reduced noise in the pH maps in **Fig. 7C** compared to **Fig. 7F**.

**Figure 7.**
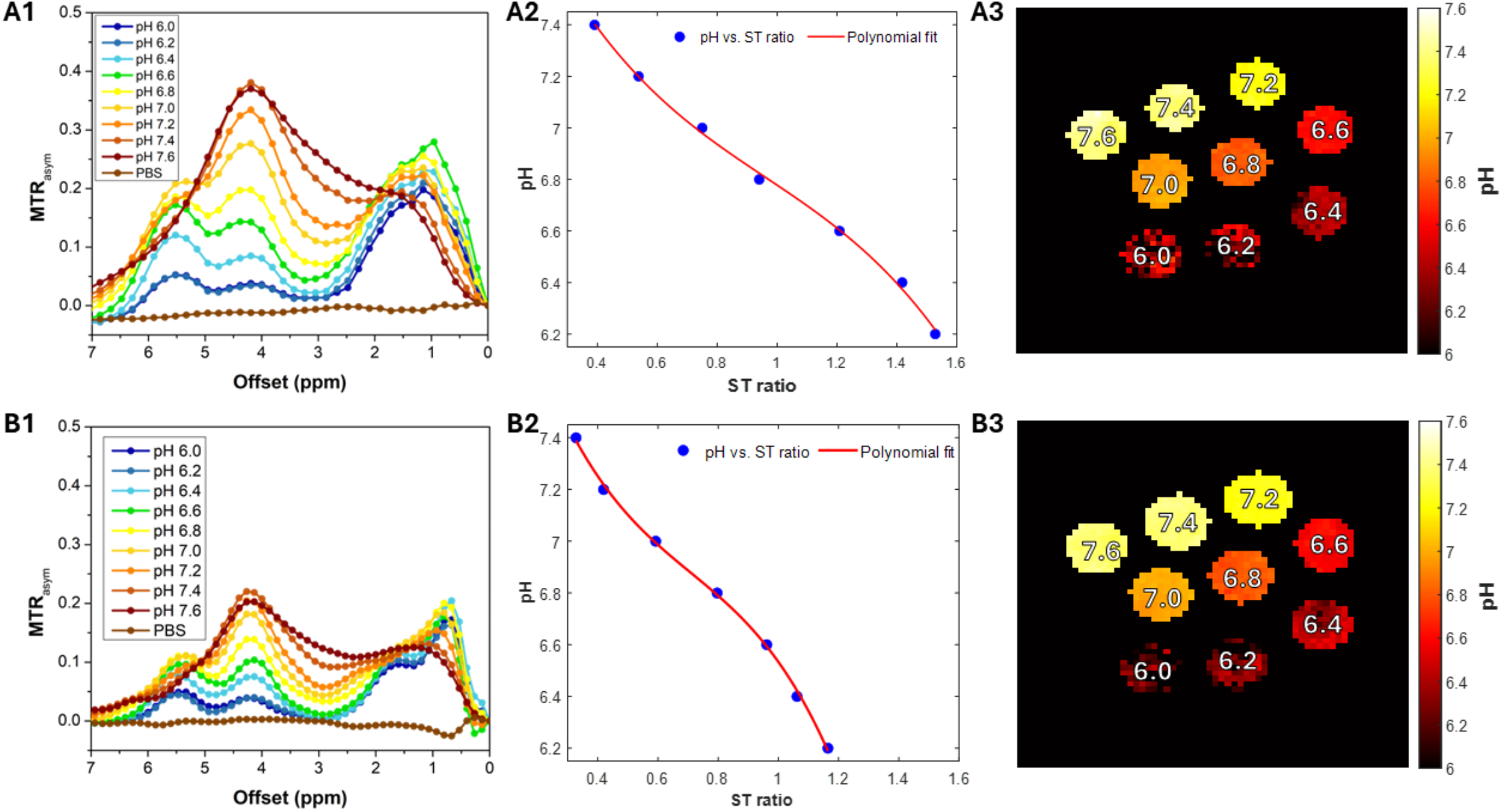
Calibration curves for pH mapping for the optimized pulse PRECISE 9.6 2500 compared to Gaussian shape. Saturation conditions: Peak B1 = 4 μT, 29 t_pul_ = 100 msec with t_mix_ = 1 msec. a) MTRasym spectra for PRECISE 9.6 2500; b) STratio curve; c) resulting phantom pH map; d) MTRasym spectra for gaussian; e) STratio curve; f) resulting phantom pH map

Finally, we decided to evaluate how the PRECISE 9.6 2500 pulse would perform for pH mapping in mice and injected 5 mice per group with 150 μL of 370 mM iopamidol. **Fig. 8** shows the results of these experiments. On comparing the two groups of mice, the CEST contrast values at 4.2 ppm and 5.5 ppm were calculated to be 5.95 ± 0.97 % and 3.80 ± 0.90 % for the PRECISE 9.6 2500 pulse and 3.49 ± 0.82 % and 3.04 ± 0.87 % for the Gaussian pulse. The peak contrast in the kidneys at 4.2 ppm was 7.8% for PRECISE 9.6 2500 as compared to 4.4% for Gaussian pulse. On performing the Mann-Whitney test, the signal enhancement for the PRECISE 9.6 2500 had a statistically significant difference compared to Gaussian. On comparing the pH values based on the ratio of signals at 4.2 and 5.5 ppm obtained for PRECISE 9.6 2500 and Gaussian, the mean pH values were found to be 6.9 ± 0.3 and 6.3 ± 0.6 respectively. The pH distribution was bell-shaped and symmetrical for PRECISE 9.6 2500 compared to Gaussian. The variance for PRECISE 9.6 2500 was 0.09 which was much lower when compared to 0.38 for Gaussian pulse. Overall, the optimized pulse significantly improved the ratiometric pH maps in live mice, similar to what was seen in phantoms.

**Figure 8.**
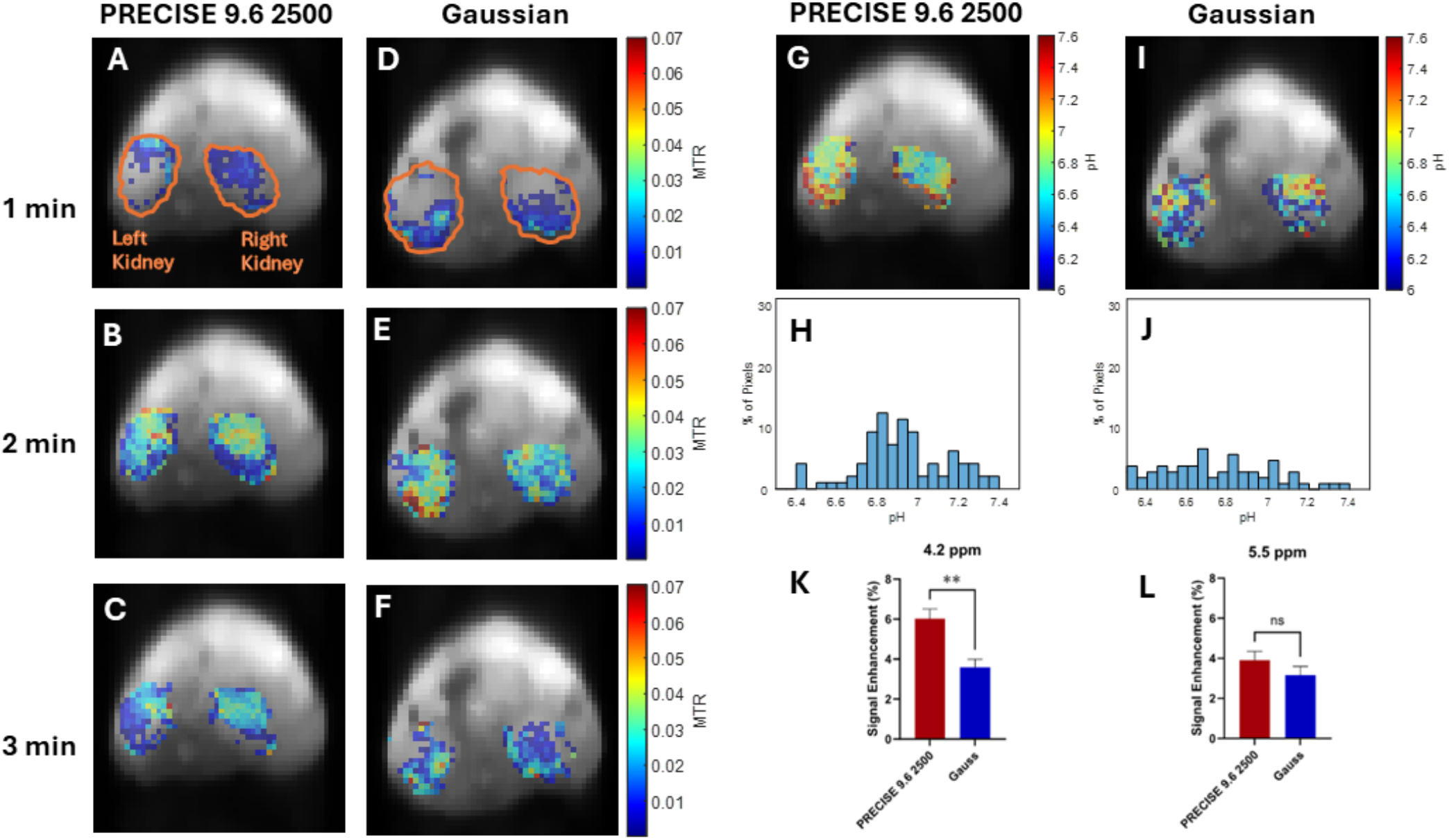
Temporal MTR_asym_ maps at 4.2 ppm obtained using (A-C) PRECISE 9.6 2500 pulse and (D-F) Gaussian pulse at 1, 2 and 3 minutes after iopamidol injection respectively. Ratiometric pH map and pH distribution histogram obtained using (G,H) PRECISE 9.6 2500 pulse and (I,J) Gaussian pulse respectively. Differences in signal enhancement and statistical significance of difference between PRECISE 9.6 2500 pulse and Gaussian pulse at (K) 4.2 ppm and (L) 5.5 ppm (n=5, P<0.05)

## Discussion

In the present study, we have focused on developing versatile pulse shapes for CEST MRI using an applied a gradient ascent algorithm to optimize the shapes for a range of Δω and k_ca_ that are observed in the agents employed in our center including the ones evaluated but also glucose^9^, superCESTide^37, 38^, creatine^39^ and others. We focused on optimizing for a 100 msec saturation pulse to be inserted into a pulse train for producing CEST MRI contrast, so these pulses are not explicitly optimized for the steady-state CEST sequences with low flip angle pulse and buildup of saturation which are also widely used on clinical scanners. Our simulations produced several candidates, and we selected 7 to test in phantoms. We compared these results to the three standard shapes with particularly excellent performance for our pulse optimized for 2,500 s^−1^, an intermediate exchange rate, using a long pulse train with short delays between the pulses. We chose 100 msec for optimizations because this was well suited for faster k_ca_, such as found in iopamidol, I45DC-diGlu, and other agents, where the shorter pulse lengths of 20, 40, or 50 msec employed in different studies are not as well suited. ^22, 25, 40^ The Fermi, sg100 and Gaussian shapes have been utilized in a number of CEST acquisitions previously. As can be seen, our best shape performed better than these on our 11.7 T scanner in phantoms using a short delay between the pulses. However, the performance will be different if lower duty cycles or steady state buildup are employed. One of the surprising findings is that the PRECISE 9.6 2500 performed even better than the PRECISE 4.2 500 pulse for the iopamidol phantom, which didn’t perform as well as the Fermi pulse either. This could be due to our choice of using the Bloch equations and a simplified 3-spin model rather than optimizing over all 4 types of exchanging spins present in iopamidol which would require 6 total spins for these simulations or it could be due to the extensive side band ripples seen in the inversion profile. The PRECISE 9.6 2500 shape performs better both in phantoms and in live mice as shown by our data. We believe this is due in large part to the higher rms integral and larger R_BW._ This waveform is quite versatile and will perform well in a wide range of circumstances based on the data we show in this manuscript.

Our study has several limitations. First, we kept the search to 100 msec with a 1 msec grid to allow for facile simulations and relatively easy-to-plug-in pulse shapes. It is worth noting that this 1 msec is shorter than the exchange lifetimes of a number of CEST agents, notably for faster k_ca_ = 4,500 s^−1^, this lifetime is a mere 0.222 msec and even for k_ca_ = 1,500 s^−1^ this lifetime is 0.67 msec. It is possible that a finer grid might improve the performance. In addition, we restricted our pulses to be equivalent across the whole train, whereas Glaser and colleagues observed in previous gradient ascent optimizations that the amplitudes might increase for ideal contrast generation over the course of the saturation module.^26^ Finally, we optimized using a relatively short total t_sat_ = 2 sec, which might not allow us to capture all the features and we employed the Bloch equations rather than using a more complete density matrix formulation. Furthermore, it could be the case that sampling over a wider range of relaxation times would be helpful, although we didn’t observe big differences between optimizations when setting T_1_ = 2 sec or 3 sec. Despite these limitations, our results indicate our optimized pulse outperforms previously employed pulse shapes.

## Conclusion

We have identified a new shape for 100 msec pulses (PRECISE 9.6 2500) that can be inserted into a wide variety of CEST pulse trains and show this generates improved CEST contrast over 3 standard shapes in both phantoms and live mice. We have also identified how this shape would change as the delay between pulses is increased. We expect this pulse to perform well in a wide range of CEST imaging sequences.

## Data Availability Statement

Pulse waveforms are available on request.

## Notes

### Competing Interest Statement

The authors have declared no competing interest.

