## Supplemental Figures for "The Proton Resonance Enhancement for CEST imaging and Shift Exchange (PRECISE) family of RF pulse shapes for Chemical Exchange Saturation Transfer MRI"

Submitted to *Magnetic Resonance in Medicine*

***Corresponding Author:**
Michael T. McMahon, Ph.D.
F.M. Kirby Research Center for Functional Brain Imaging
Kennedy Krieger Institute
707 N. Broadway Ave.
Baltimore, MD, 21205


Grant support from NIH: R01DK121847, R01EB030565
Running title: Optimized pulses for CEST MRI

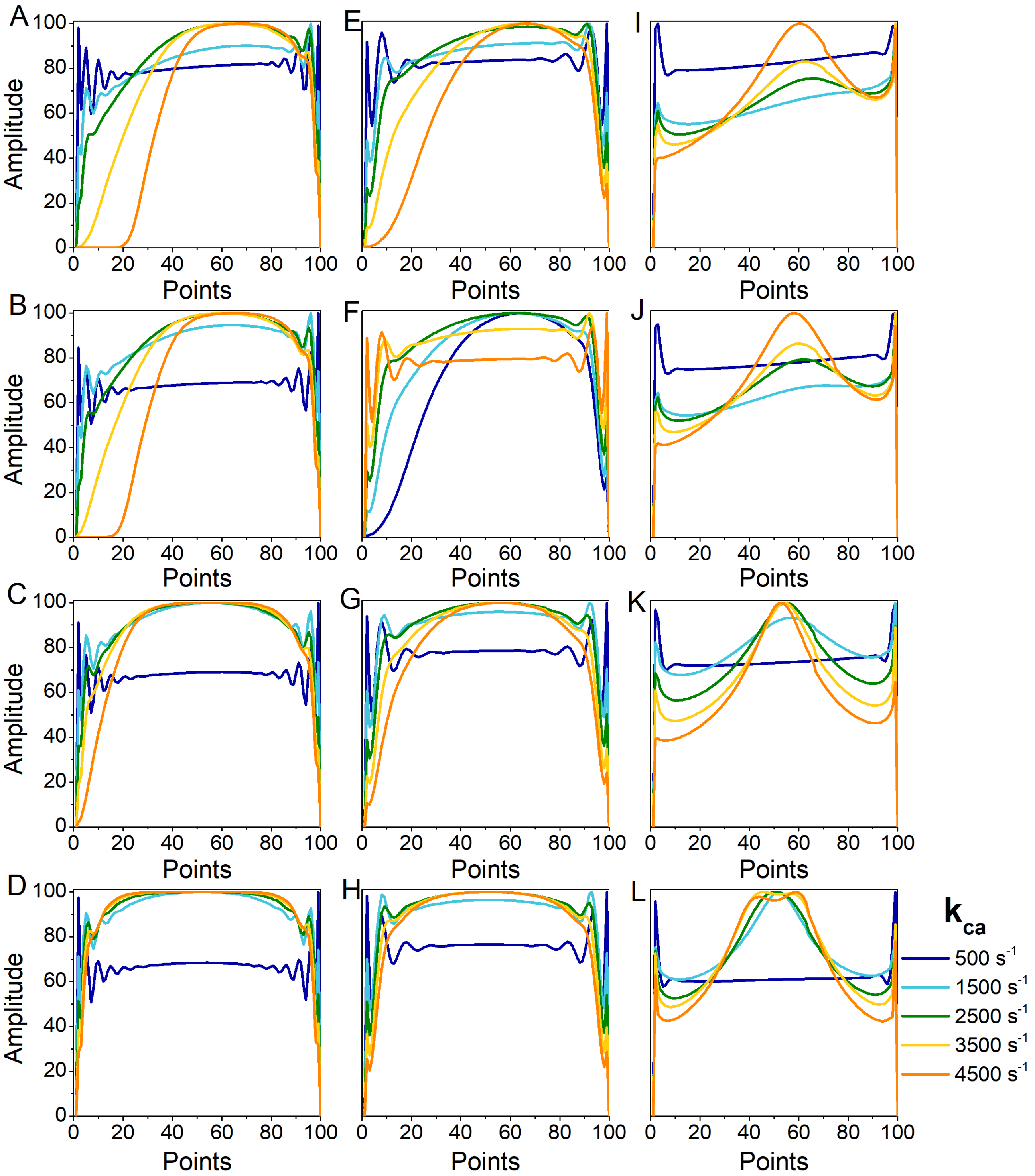


**Figure S1.** Optimized pulse shapes as a function of exchange rate and offset. 9.6 ppm pulse shapes including $\tau_{mix}$ = A) 1 msec; B) 10 msec; C) 40 msec; D) 100 msec. 4.2 ppm pulse shapes including: $\tau_{mix}$ = D) 1 msec; E) 10 msec; F) 40 msec; G) 100 msec. 2.0 ppm pulse shapes including $\tau_{mix}$ = H) 1 msec; I) 10 msec; J) 40 msec; K) 100 msec.

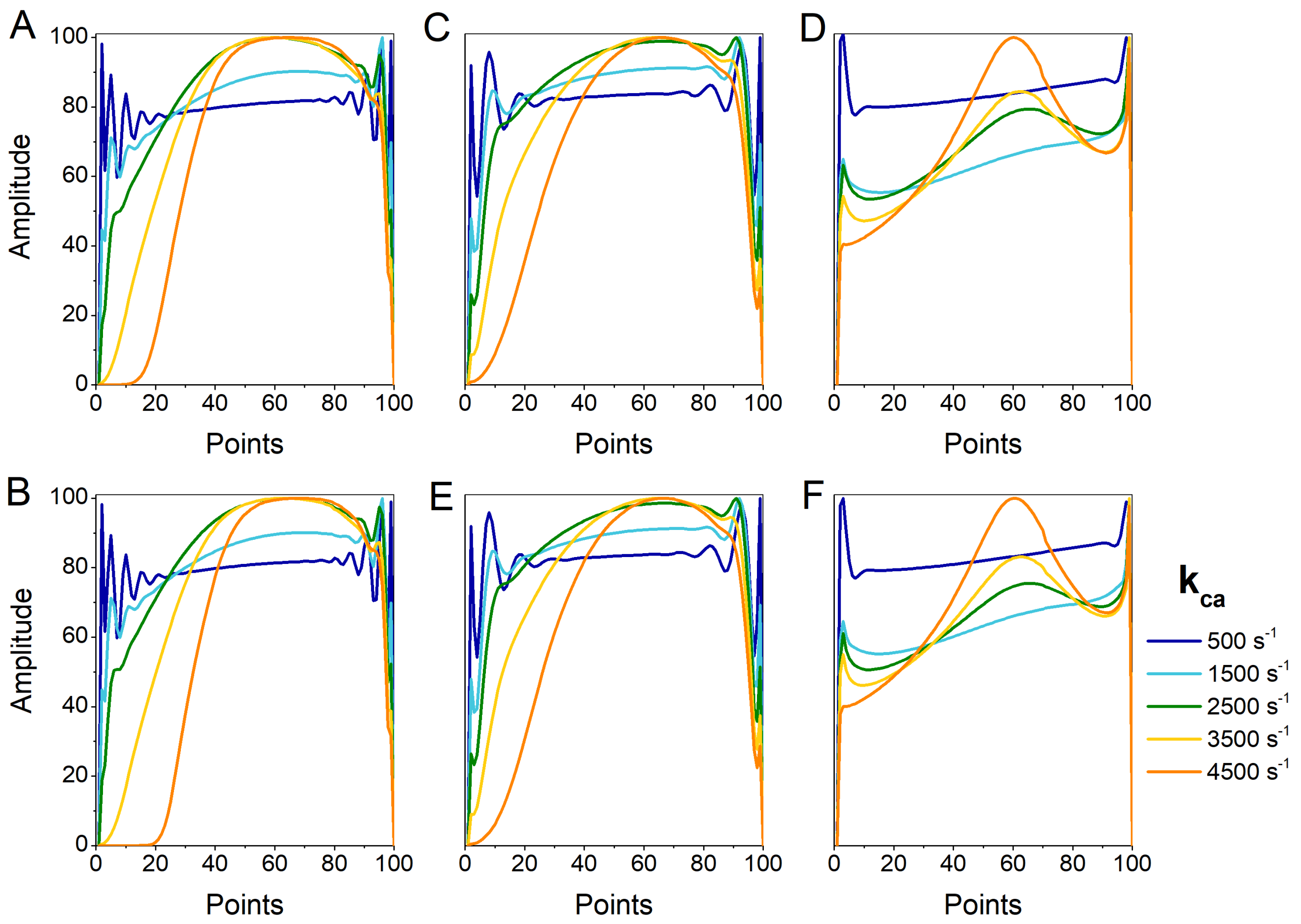


**Figure S2.** Dependence of optimized pulse shape with water $T_1$ for A) offset = 9.6 ppm, $T_{1a}$ = 2 sec; B) offset = 9.6 ppm, $T_{1a}$ = 3 sec; C) offset = 4.2 ppm, $T_1$a = 2 sec; D) offset = 4.2 ppm, $T_{1a}$ = 3 sec; E) 2 ppm, $T_{1a}$ = 2 sec; F) offset = 2 ppm, $T_{1a}$ = 3 sec;
